# The inhaled corticosteroid ciclesonide blocks coronavirus RNA replication by targeting viral NSP15

**DOI:** 10.1101/2020.03.11.987016

**Authors:** Shutoku Matsuyama, Miyuki Kawase, Naganori Nao, Kazuya Shirato, Makoto Ujike, Wataru Kamitani, Masayuki Shimojima, Shuetsu Fukushi

## Abstract

Steroid compounds, which are expected to have dual functions in blocking host inflammation and MERS-CoV replication, were screened from a chemical library. Within this library, ciclesonide, an inhaled corticosteroid, suppressed human coronavirus replication in cultured cells, but did not suppress replication of respiratory syncytial virus or influenza virus. The effective concentration of ciclesonide to block SARS-CoV-2 (the cause of COVID-19) replication (EC_90_) was 6.3 μM. After the eleventh consecutive MERS-CoV passage in the presence of ciclesonide, a resistant mutation was generated, which resulted in an amino acid substitution (A25V) in nonstructural protein (NSP) 15, as identified using reverse genetics. A recombinant virus with the mutation was also resistant to ciclesonide suppression of viral replication. These observations suggest that the effect of ciclesonide was specific to coronavirus, suggesting this is a candidate drug for treatment of patients suffering MERS or COVID-19.

## Introduction

The COVID-19 outbreak began on December 2019 in Wuhan, China (1), and the causative virus, SARS-CoV-2, has rapidly spread into 43 countries as of February 27, 2020. The number of infected patients exceeds 81,000, and the death toll exceeds 2,700. Several drugs, including lopinavir, remdesivir, and chloroquine, have been reported to be presumably effective in treating this disease (2, 3).

Systemic treatment with corticosteroids is contraindicated for the severe pneumonia caused by viruses such as MERS-CoV and SARS-CoV, as steroids suppress the innate immune system, resulting in increased viral replication. In fact, for SARS in 2003 and MERS in 2013, treatment with corticosteroids was associated with increased mortality (4, 5). In the present study, we reconsider the use of inhaled corticosteroids, which have been excluded from the treatment of pneumonia caused by coronavirus.

## Results and Discussion

Steroid compounds, which are expected to have dual functions in blocking both coronavirus replication and host inflammation, were screened from a chemical library. The cytopathic effect caused by MERS-CoV infection was measured to evaluate viral replication. Four steroid compounds, ciclesonide, mometasone furoate, mifepristone, and algestone acetophenide conferred a greater than 95% cell survival rate (Fig. S1).

Next, concentration-dependent viral growth suppression and drug cytotoxicity were assessed. Ciclesonide exhibited low cytotoxicity and potent suppression of viral growth (Fig. 1a). Cortisone and prednisolone, which are commonly used for systemic steroid treatment, dexamethasone, which has strong immunosuppressant effects, and fluticasone, a commonly used inhaled steroid, did not suppress viral growth (Fig. 1a).

**Figure 1:**
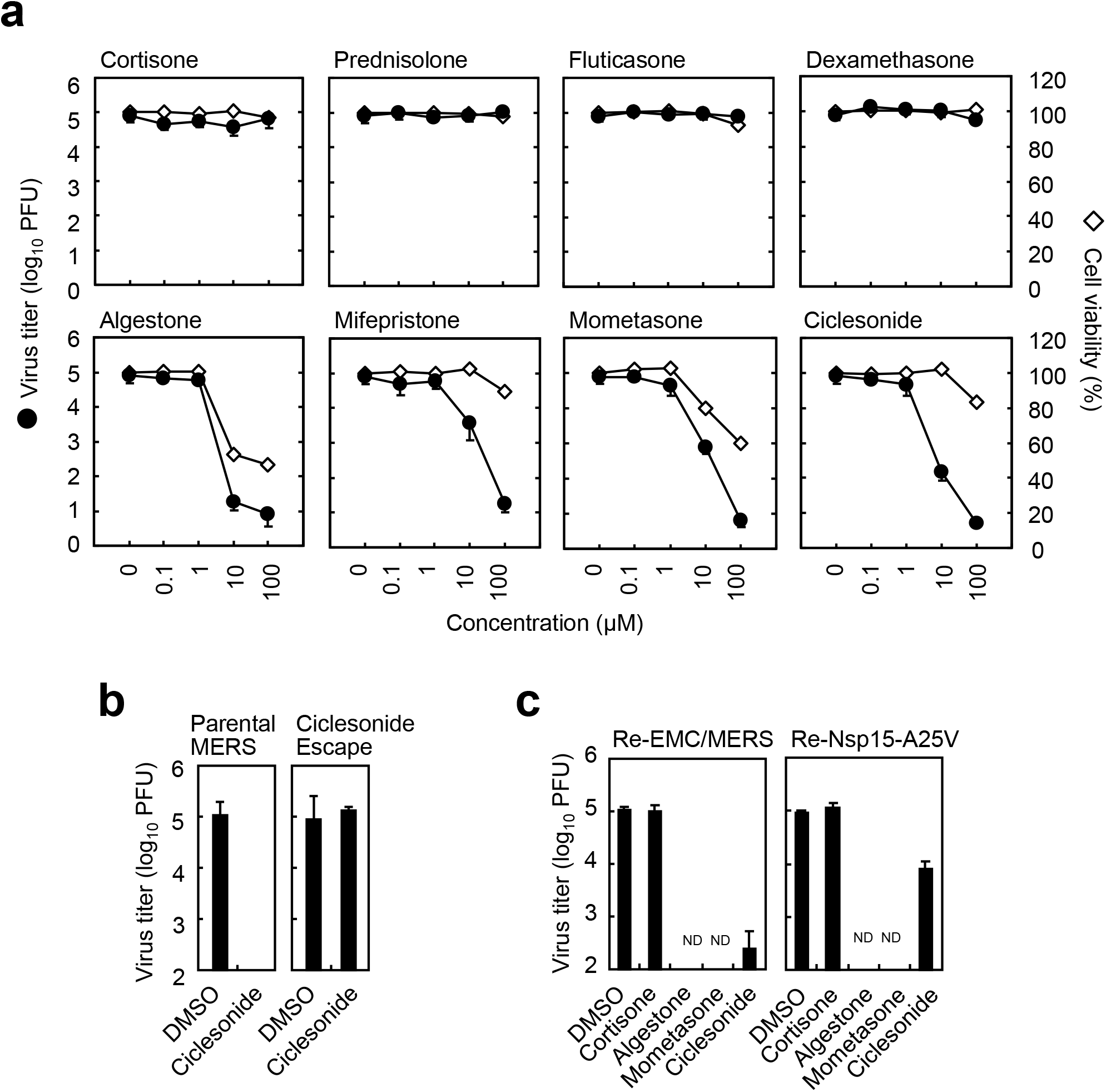
Antiviral effects of steroids on MERS-CoV infection. **(a)** Effect of steroids on MERS-CoV replication. Vero cells were infected with MERS-CoV at MOI = 0.01 in the presence of steroids for 24 h. The viral titer in the cell supernatant was quantified by standard plaque assay using Vero/*TMPRSS2* cells. Cell viability in the presence of steroids was quantified by WST assay at 24 hours post-infection (hpi). **(b)** Ciclesonide escape mutant. Vero cells treated with ciclesonide were infected with parental MERS-CoV or ciclesonide escape mutant. Viral titer was measured as described in panel a. **(c)** Vero cells were infected with the parental MERS-CoV/EMC strain (Re-EMC/MERS) or the recombinant mutant strain (Re-Nsp15-A25V) with an amino acid substitution at A25V in NSP15. Viral titer was measured as described in panel a.

The antiviral effects of steroids against various species of viruses were tested by quantifying propagated virus in culture medium. Ciclesonide and mometasone also suppressed replication of other coronaviruses, HCoV-229E and SARS-CoV, but not replication of RS virus or influenza virus (Fig. S2). In addition, ciclesonide slightly but significantly inhibited rubella virus (having positive strand RNA genome) replication (Fig. S2), suggesting that ciclesonide interacted with the replication site of positive-strand RNA virus intracellularly.

To identify the drug target of virus replication, we conducted 11 consecutive MERS-CoV passages in the presence of 40 μM ciclesonide or 40 μM mometasone. A mutant virus that developed resistance ciclesonide was generated (Fig. 1b), but no resistant virus to mometasone was generated. Next-generation sequencing identified that an amino acid substitution at A25V (C19647T in the reference sequence NC_019843.3) in nonstructural protein 15 (NSP15), coronavirus endoribonuclease (6–8), was the predicted mechanism for viral resistance to ciclesonide. A recombinant virus carrying the A25V amino acid substitution (Re-Nsp15-A25V) was generated from the parental MERS-CoV/EMC strain (Re-EMC/MERS) with a BAC reverse genetics system (9), which overcame the antiviral effect of ciclesonide (Fig. 1c). Interestingly, the mutant virus was inhibited by mometasone, suggesting that the antiviral target of mometasone was different from that of ciclesonide.

Next, the effect of ciclesonide on SARS-CoV-2 infection was evaluated. VeroE6/*TMPRSS2* cells(10) were infected with authentic SARS-CoV-2 in the presence of steroids or other inhibitors. At 6 h post-infection, cellular RNA was isolated, and real-time PCR was conducted to quantify the amount of viral RNA (11). Ciclesonide and mometasone suppressed viral replication with a similar efficacy to lopinavir (Fig. 2a). For comparison, viral cell entry inhibitors were tested. E64d substantially reduced viral RNA levels, but nafamostat and camostat had only modest antiviral effects, suggesting that SARS-CoV-2 primarily utilized the cathepsin/endosomal pathway of cell entry rather than the TMPRSS2/cell surface pathway to enter TMPRSS2-expressing cells, consistent with a recent study (12). Ciclesonide blocked SARS-CoV-2 replication at low concentrations (EC_50_ =□4.4□μM; EC_90_□=□6.3□μM; CC_50_□>□100□μM) (Fig. 2b).

**Figure 2:**
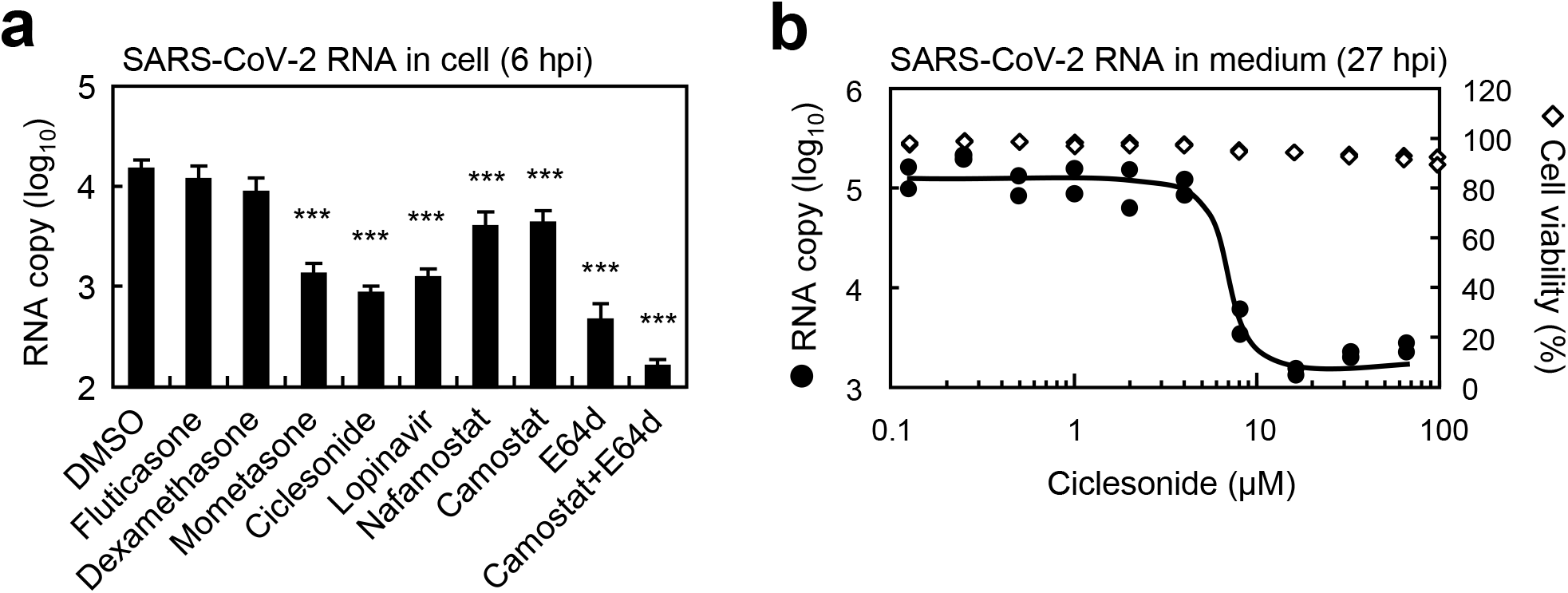
Antiviral effects of steroids on SARS-CoV-2. **(a)** Intracellular SARS-CoV-2 RNA (6 hpi). VeroE6/*TMPRSS2* cells were infected with SARS-CoV-2 at MOI = 1 in the presence of 10□μM compounds for 6 h. Cellular viral RNA was quantified by real-time PCR using the E gene primer/probe set. **(b)** Culture Medium SARS-CoV-2 RNA (27 hpi). VeroE6/*TMPRSS2* cells were infected with SARS-CoV-2 at MOI = 0.01 in the presence of ciclesonide for 27 h. Viral RNA in culture medium was quantified by real-time PCR using the E gene primer/probe set. Cell viability in the presence of ciclesonide was quantified at 27 hpi by WST assay.

This study suggests that ciclesonide presumably interacts with viral NSP15, either directly or indirectly, to suppress viral replication of SARS-CoV-2. An amino acid substitution in the ciclesonide-resistant mutant, located at the dimerization site of the NSP15 homo-hexamer (12), suggested that ciclesonide interacts with NSP15 during viral biogenesis. Future studies will provide a detailed analysis of the molecular mechanisms of ciclesonide suppression of viral replication.

Ciclesonide is a safe drug, and can be administered to infants at high concentrations. Inhaled ciclesonide is expected to reduce viral replication and host inflammation in the lungs, with decreased immunosuppressive effects compared to systemic corticosteroids, as ciclesonide primarily remains in the lung tissue, and does not significantly enter the bloodstream. Treatment of patients with ciclesonide should be carefully evaluated, considering the benefit-risk balance of the drug.

## Materials and Methods

### Cells and viruses

Vero cells and VeroE6 cells expressing *TMPRSS2* (VeroE6/*TMPRSS2*) were maintained in Dulbecco’s modified Eagle medium (DMEM; Sigma-Aldrich, USA) supplemented with 5% fetal bovine serum (Gibco-BRL, USA). MERS-CoV and SARS-CoV-2 were propagated in Vero and VeroE6/*TMPRSS2* cells.

### Steroids and inhibitors

The following compounds were used: cortisone, prednisolone, fluticasone, dexamethasone, algestone acetophenide, mifepristone, mometasone furoate, and ciclesonide (from Prestwick the Chemical Library; PerkinElmer, MA, USA), nafamostat (N0289; Sigma), camostat (3193; Tocris Bioscience, USA), E64d (330005; Calbiochem, USA), and lopinavir (SML1222; Sigma).

### Quantification of coronaviral RNA

Confluent cells in 96-well plates were inoculated with virus in the presence of steroid compounds. Cellular RNA was isolated at 6 hours post-infection (hpi) using a CellAmp Direct RNA Prep Kit (3732; Takara, Japan) and culture medium collected at 24 hpi or 27 hpi was diluted 10-fold in water, then boiled. A real-time PCR assay was performed to quantify the amount of coronavirus RNA with a MyGo Pro instrument (IT-IS Life Science, Ireland) using primers and probes described previously (11, 13).

### Cytotoxicity Assays

Confluent cells in 96-well plates were treated with steroids. After incubation for 24 or 27 hours, a cell viability assay was performed using WST reagent (CK12; Dojin Lab, Japan) according to manufacturer’s instructions.

### Generation of recombinant MERS-CoV from BAC plasmids

A bacterial artificial chromosome (BAC) clone carrying the full-length infectious genome of the MERS-CoV EMC2012 strain was used to generate recombinant MERS-CoV, as described previously (9).

### Statistical analysis

Statistical significance was assessed using an ANOVA test. A P-value < 0.05 was considered statistically significant. In figures, significance is indicated as follows: n.s., not significant; * P ≤ 0.05; ** P ≤ 0.01; and *** P ≤ 0.001. Error bars indicate standard deviations (SD).

## Acknowledgements

We thank Tsuneo Morishima of Aichi Medical University for helpful suggestions. We also thank Ron A. M. Fouchier and Bart L. Haagmans of Erasmus Medical Center for providing MERS-CoV, and John Ziebuhr of the University of Wurzburg for providing SARS-CoV. This study was supported by Grants-in Aid from the Japan Agency for Medical Research and Development (AMED) (Grant Number JP19fk0108058j0802), and from the Japan Society for the Promotion of Science (JSPS) (Grant Number 17K08868).

## Author contributions

S.M., M.S., and S.F. designed the research; S.F. screened drugs; S.M., M.K., K.S., and M.U. performed experiments; N.N. performed NGS; W.K. performed reverse genetics; S.M. wrote the paper.

The authors declare no competing interests.

