## Supplementary material for "The inhaled corticosteroid ciclesonide blocks coronavirus RNA replication by targeting viral NSP15": Suppl Figs

**Abstract**

Steroid compounds, which are expected to have dual functions in blocking host inflammation and MERS-CoV replication, were screened from a chemical library. Within this library, ciclesonide, an inhaled corticosteroid, suppressed human coronavirus replication in cultured cells, but did not suppress replication of respiratory syncytial virus or influenza virus. The effective concentration of ciclesonide to block SARS-CoV-2 (the cause of COVID-19) replication (EC_90_) was 6.3 μM. After the eleventh consecutive MERS-CoV passage in the presence of ciclesonide, a resistant mutation was generated, which resulted in an amino acid substitution (A25V) in nonstructural protein (NSP) 15, as identified using reverse genetics. A recombinant virus with the mutation was also resistant to ciclesonide suppression of viral replication. These observations suggest that the effect of ciclesonide was specific to coronavirus, suggesting this is a candidate drug for treatment of patients suffering MERS or COVID-19.

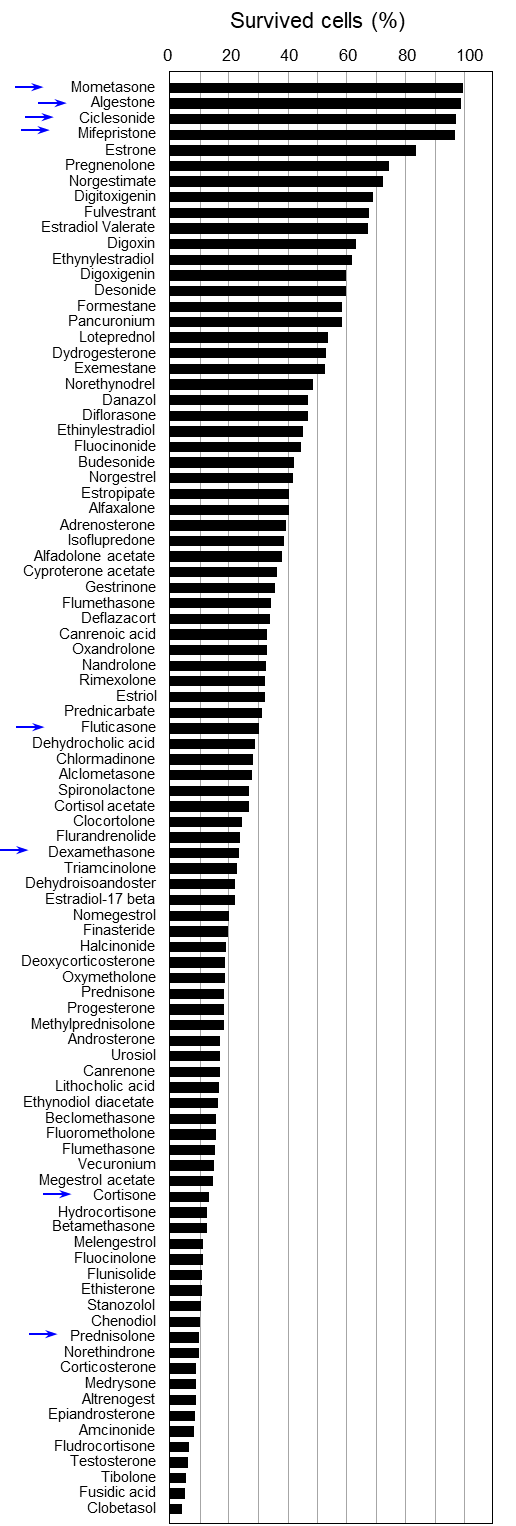

**Figure S1: Screening of steroid compounds on MERS-CoV viral replication**. Vero cells seeded on 96-well microplates were infected with 100 TCID_50_ MERS-CoV in the presence of 92 steroid compounds (10 μM) from the Prestwick the Chemical Library (PerkinElmer), and the cytopathic effect was observed at 72 hpi. Surviving cells were stained with crystal violet and subsequently photographed and quantified with ImageJ software. Arrows indicate the steroid compounds further assessed in this study.

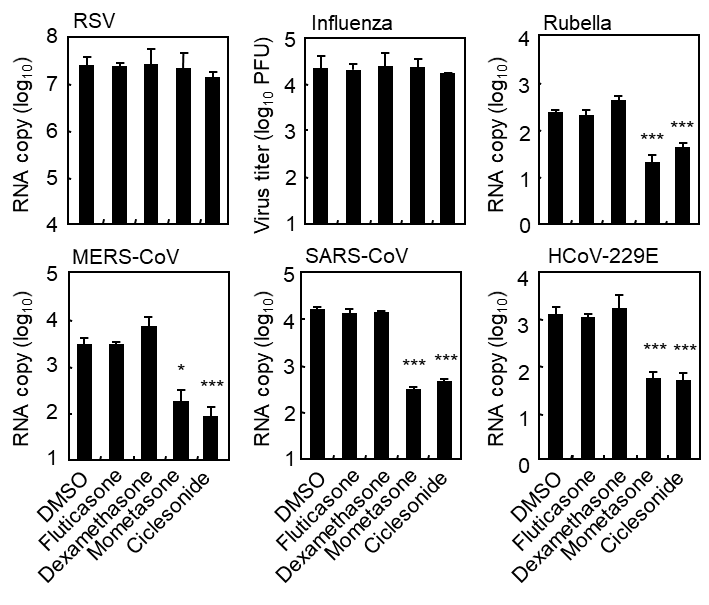

**Figure S2: Antiviral effects of steroid compounds on other viral species.** Cells were infected with viruses at MOI = 0.01 in the presence of steroids. Cell supernatant viral yield was quantified by standard plaque assay or real-time PCR methods recommended by the US CDC. Virus, cells, and incubation period were as follows: Respiratory syncytial virus A (RSV-A long) to Hep2 cells for 1 day, Influenza H3 (CA04) to MCDCK cells for 1 day, Rubella virus (TO336) to Vero cells for 7 days, MERS-CoV (EMC) to Vero cells for 1 day, SARS-CoV (Frankfurt-1) to Vero cells for 1 day, and HCoV-229E (VR-740) to HeLa cells for 1 day. Error bars indicate the SD of the means from four independent wells. significance is indicated as follows: n.s., not significant; * P ≤ 0.05; ** P ≤ 0.01; and *** P ≤ 0.001. Error bars indicate standard deviations (SD).
